# Wildlife as a sentinel for pathogen introduction in nonendemic areas: first detection of *Leishmania tropica* in wildlife in Spain

**DOI:** 10.1101/2024.03.16.585353

**Authors:** Iris Azami-Conesa, Pablo Matas Méndez, Paula Pérez-Moreno, Javier Carrión, J.M. Alunda, Marta Mateo Barrientos, María Teresa Gómez-Muñoz

## Abstract

Leishmaniasis is a chronic global arthropod-borne zoonotic disease produced by several species of *Leishmania*, with cutaneous, mucocutaneous, and visceral clinical manifestations. In Spain, only *Leishmania infantum* has been reported so far, although other species of *Leishmania*, such as *L. tropica* and *L. major*, are present in surrounding countries.

The aim of this work is to analyze the occurrence of *Leishmania* spp. infection in European wildcats (*Felis silvestris*) as sentinels, including their genotypic characterization. Necropsies of 18 road killed wildcats were conducted. Samples of ear skin and spleen were taken for DNA isolation and PCR of the highly sensitive *SSUrDNA* target. Subsequent PCR tests were performed using more specific targets for the determination of *Leishmania* species: *hsp70* and *ITS1*. Positive samples were sequenced, and phylogenetic trees constructed. Seven wildcats were found positive for *Leishmania* spp.. Based on the *hsp70* and *ITS1* sequences, an animal was found to be infected only with *L. tropica* in ear skin samples, while two cats were found to be infected with *L. infantum* in both the ear skin and the spleen. In one animal, a clear sequence of *L. infantum* ITS1 and a sequence of *L. tropica hsp70* were obtained from the ear skin. Since hsp70 and ITS1 sequencing was not possible in three cats, the species of *Leishmania* infecting them was not determined.

This is the first report of autochthonous infection with *L. tropica* in the Iberian Peninsula. Health care professionals, including physicians, dermatologists, and veterinarians, must be aware of this for a correct diagnosis, treatment, and management of possible co-infections.

## Introduction

Leishmaniasis is a global zoonotic disease transmitted by the bite of phlebotomine insects, with more than 1,000,000 new cases of the cutaneous form and 30,000 new cases of the visceral form estimated every year, in 92 endemic countries [1]. The clinical presentation in humans is variable and includes cutaneous (CL), mucocutaneous (MCL), visceral (VL), and post Kala-azar dermal leishmaniasis (PKDL). The disease is caused by several species of *Leishmania*, with the *L. donovani* complex, the *L. major* and the *L. tropica* complex being the most frequent in Africa, Asia, southern Europe, South America, and the Middle East, while the *L. mexicana* complex, the *L. guyanensis* complex and the *L. braziliensis* complex are the most diagnosed in the Americas. The cutaneous form of the disease is the most common and is responsible for the stigmatization and social isolation of thousands of people, particularly women, while VL is usually lethal without treatment and is caused by *L. donovani* in Africa and Europe, and *L. infantum* in southern Europe, west and central Asia, and the Americas. MCL is characterized by the destruction of mucous tissue of the nose, palate, and pharynx, and can severely affect the daily life of thousands of people [2,3]. Human leishmaniasis is more prevalent in low-income countries with displaced populations due to war, social, or economic conflicts, and is classified by the WHO as a neglected tropical disease (NTD). This scenario makes early diagnosis and treatment of the disease, as well as the introduction of preventive measures, difficult [3].

In Europe, VL is caused by *L. infantum*, with humans and dogs as the target species. The infection is more prevalent in southern countries such as Portugal, Spain, Italy, and Greece. Dogs are heavily affected by this zoonotic species [4] and are considered the main reservoir. Human infections mainly affect immunocompromised patients (such as HIV + or recipients of solid organ transplants), as well as occasional reports in elderly and children [5]. Several studies have stressed the paramount importance of wildlife in maintaining *Leishmania* infection [6,7]. Wildcats (*Felis silvestris*) are susceptible to the same pathogens as domestic cats and therefore could be potential reservoirs for the infection. However, the studies conducted have been scarce [8–10], with a limited number of animals analysed in each study (n=4, n=3, n=4, respectively). The three studies identified *L. infantum* using a small fragment of kDNA or qPCR from the ITS2 regions.

Despite suitable bioclimatic conditions and vectors available for anthroponotic *L. tropica* (*Phlebotomus sergenti*) and *L. major* (*P. papatasi*) [11,12, only *L. infantum* has been reported in the Iberian Peninsula [6,12,13]. However, *L. major* has recently been reported in a cat from Lisbon, Portugal [14]. Moreover, it is worth noting that *L. tropica* has been previously reported in Greece [15] and the three species (*L. infantum, L. tropica*, and *L. major*) are present in Morocco [16], as well as in other countries bordering Europe [17]. The availability of a larger number of wildcats allowed us to identify the presence of *Leishmania* in various tissues and genetically characterize the samples. The aim of this work is to analyze the occurrence of *Leishmania* spp. infection in wildcats as sentinels, including their genotypic characterization.

## Materials and Methods

### Animals, samples, and geographic area

In this study, samples of 18 European wildcats (*Felis silvestris*) that were killed on the road during 2022 and 2023 in the Spanish provinces of Ávila, Burgos, Ciudad Real, Guadalajara, Soria, and Valladolid,,, were analyzed. Samples from the skin and spleen were selected, since they are the preferred sites for cutaneous and/or visceral *Leishmania* amastigotes. All roadkilled wildcats were included in the study period. Animals were transported to the wildlife recovery centers of Castilla y León (Burgos, Valladolid) and Castilla-La Mancha (Ciudad Real, Guadalajara), systematic necropsies were performed, and the presence/absence of lesions annotated. A portion of the ear skin and spleen was taken and frozen at -20°C for further processing. The dates, age, sex, and location (geographic coordinates where the dead animals were found) were included in the data base.

### Maps

The map shown in figure 1 was created taking as reference the European Centre for Disease Prevention and Control (ECDC) maps of distribution in Europe of *Phlebotomus sergenti*, the main vector of *L. tropica*, in 2022 and 2023 [18], and the Universal Transverse Mercato (U.T.M.) geographic coordinates of the closest villages to where the wildcats analysed in this study were found. The AutoCAD program (Autodesk® Inc., Landmark, San Francisco, CA, USA) was used to observe the overlaps between the two maps.

**Figure 1.**
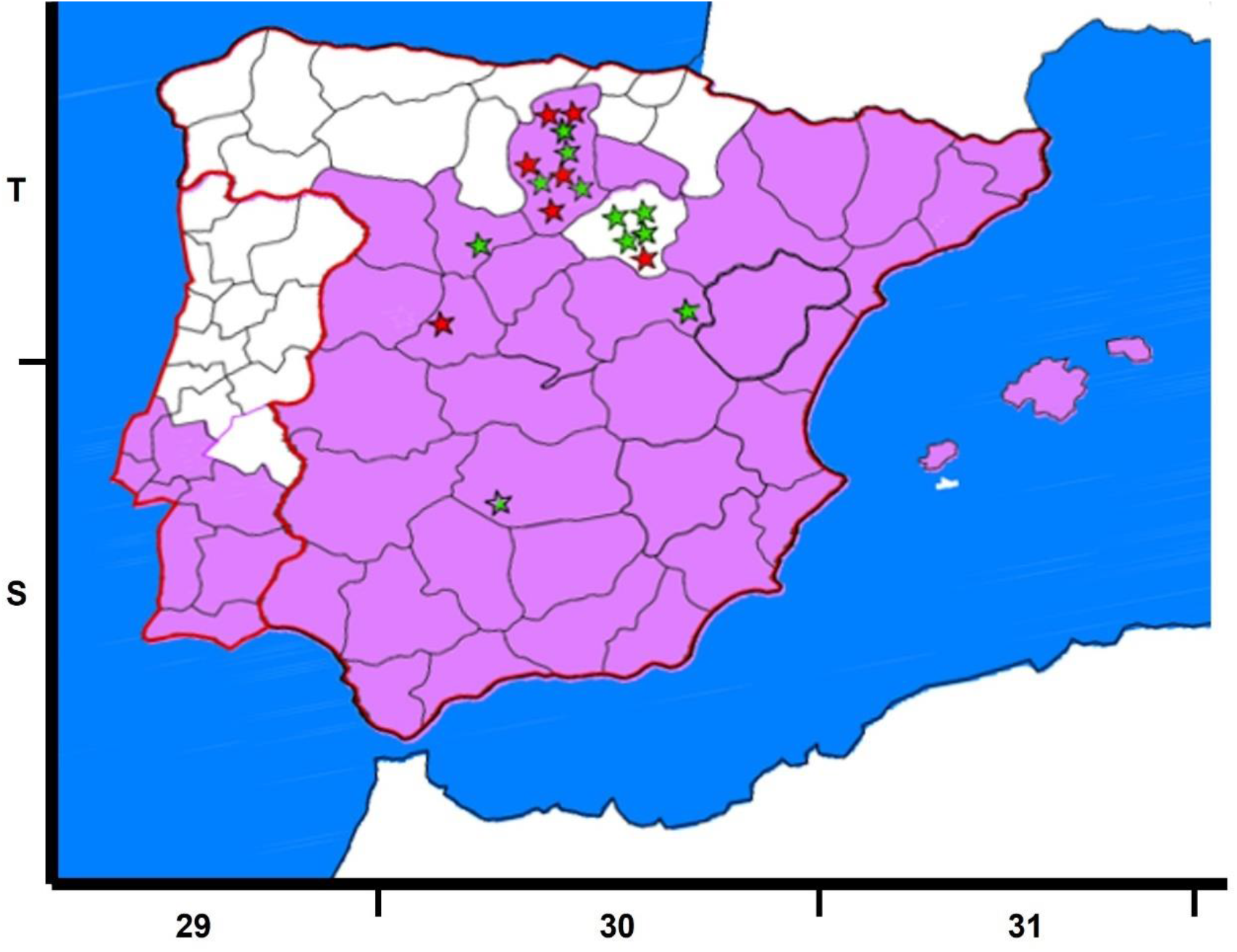
Map of the Iberian Peninsula with U.T.M. geographic coordinates. The purple area corresponds to the distribution of *Phlebotomus sergenti* in 2022 and 2023 available at ECDC [18]. Red stars indicate the detection and identification of *Leishmania* spp. by *SSU-rDNA sequences*. Green stars indicate the absence of *Leishmania* spp. in the sampled cats.

### DNA isolation

DNA was isolated from the spleen (10 mg) and the skin of the ears (20 mg) using the NZYTech Tissue gDNA isolation kit (NZYTech, Portugal) and following the manufacturer’s instructions. DNA elution was performed in 60 µL of elution buffer instead of 100 µL to increase DNA concentration. Positive and negative controls were used in each batch of the experiment. The positive control consisted of samples derived from *L. major* (clone V1: MHOM/IL/80/Friedlin) or *L. braziliensis* (MHOM/BR/75/M2904) promastigotes in culture (1×10^6^ promastigotes), while the negative control consisted of water free of DNase. The DNA content was measured on a Multiskan GO microplate spectrophotometer with µDrop™ plates (Thermo Fisher Scientific, MA, USA). DNA from cultures of *L. major* and *L. braziliensis* was used as internal controls for sequencing also. These two isolates were chosen because they were absent from Spain.

### PCR of small subunit ribosomal RNA (SSU-rDNA), Internal Transcribed Spacers-5.8 ribosomal DNA 1 (ITS1), and heat shock protein 70 (hsp70) targets

DNA samples from the skin of the ear and spleen of the 18 cats were used for *Leishmania* DNA detection. DNA from different species of *Leishmania* was also used as an internal control for PCR and sequencing; *L. major* (clone V1: MHOM/IL/80/Friedlin) and *L. braziliensis* (MHOM/BR/75/M2904). Only samples with good DNA quality (absorbance ratio at 260/280 nm > 1.7) were used for all PCRs. All PCR conditions have been previously described and adapted to the working conditions recommended by the enzyme manufacturer (Supreme NZYTaq II 2x Master Mix, NZYTech, Lisbon, Portugal).

*The SSU-rDNA* target was chosen as the first PCR because of its higher sensitivity compared to other PCR targets. Nested PCRs (nPCR) to amplify a 358 bp fragment of the *SSU-rDNA* target were performed using primers R221 (5′-GGTTCCTTTCCTGATTTACG-3′) and R332 (5′-GGCCGGTAAAGGCCGAATAG-3′) for the outer section, and primers R223 (5′-TCCATCGCAACCTCGGTT-3′) and R333 (5′-AAGCGGGCGCGGTGCTG-3′) for the inner section [19], following previously described conditions [20].

Primers L5.8S (5′-ACACTCAGGTCTGTAAAC-3′) and LITSR (5′-CTGGATCATTTTCCGATG-3′) were used for the outer section of the nPCR for *ITS1* region and primers SAC (5′ - CATTTTCCGATGATTACACC-3′) and VAN2 (5′-CGTTCTTCAACGAAATAGG-3′) were used for the inner section to amplify a fragment of 280-330 bp, which allows species determination, as previously described [20,21].

The nPCR used for *hsp70* amplified a fragment of 741 bp and was based on the primers and methods previously described for phylogenetic purposes [14,22]. Briefly, the external primers HSP70-F25 (5′-GGACGCCGGCACGATTKCT-3′) and HSP70-R1310 (5′-CCTGGTTGTTGTTCAGCCACTC-3′) were used at 0.8 µM each, and the internal primers HSPF (5′-GACAACCGCCTCGTCACGTTC-3′) and HSPR (5′-GTCGAACGTCACCTCGATCTGC-3′) were used at 0.4 µM each. In the first PCR, 40 cycles of 94 ° C for 40 s, 61 ° C for 60 s and 72 ° C for 120 s were used, while in the second PCR, 40 cycles of 94 ° C for 40 s, 61 ° C for 60 s and 72 ° C for 60 s were used.

All PCR reactions were performed in 25 µL using 12.5 µL of Supreme NZYTaq II 2x Master Mix. An initial step of 95 ° C for 5 min to activate the enzyme and a final elongation step of 72 ° C for 10 min to allow the elongation of the PCR products were used in all PCR reactions. In the first PCR, 5 µL of DNA was used, while in the second PCR reactions 5 µL of a 1:50 dilution of each first PCR product were used for *hsp70*, while a 1:40 dilution of the first PCR product was used for *SSU-rDNA* and *ITS1*. The results of nPCR were visualized on 1% agarose gels stained with SYBR™ safe DNA gel staining (Invitrogen, Thermo Fisher Scientific, MA, USA) under UV light.

### Sequencing and alignment

Samples containing the amplicons of the expected size were sent to the facilities of Macrogen Spain (Madrid, Spain) for Sanger sequencing in forward and reverse directions. The resulting sequences were aligned using Molecular Evolutionary Genetics Analysis v.11 software (MEGA XI) [23] and FinchTV 1.4.0 software (Geospiza, Inc.; Seattle, WA, USA), manually checked, and subjected to BLAST analysis using the nucleotide Basic Local Alignment Search Tool (BLAST) (National Library of Medicine, Rockville, MD, USA). Only sequences with more than 150 bp that were clear in both directions were sent to GenBank for accession numbers. PCR and sequencing were repeated at least twice for each sample to ensure results.

### Phylogenetic trees

Phylogenetic trees were constructed using the sequences obtained in the present study and other sequences available from GenBank. Only the trees for *hsp70* and *ITS1* are shown because the *SSU-rDNA* target is a conserved region and only a difference of 1-2 bp was observed between the sequences obtained. In all trees, a sequence from *Trypanosoma cruzi* was included as an outgroup reference. Additionally, other sequences of *Leishmania* spp. from GenBank and sequences from *L. infantum* and the species of *Leishmania* mentioned above were included from cultures to verify similarity (*L. major* and *L. braziliensis*). The sequences had a minimum length of 292 nucleotides for the *ITS1* region and 635 nucleotides for *hsp70*. Evolutionary history was inferred using the maximum likelihood method based on the Tamura–Nei model [24]. The tree with the highest logarithmic likelihood was displayed. The percentage of trees in which the associated taxa clustered together was shown next to the branches. Initial trees for the heuristic search were obtained automatically by applying the neighbor-joining and BioNJ algorithms to a pairwise distance matrix estimated using the maximum composite likelihood (MCL) approach and selecting the topology with the superior log likelihood value. The trees were drawn to scale, and the lengths of the branches were measured on the basis of the number of substitutions per site. The analysis included 13-18 nucleotide sequences, depending on the tree. The included codon positions were 1st + 2nd + 3rd + Noncoding. All positions that contained gaps and missing data were eliminated. Evolutionary analyzes were conducted via MEGA XI [23]. A bootstrap of 2000 replicates was used to assess the reliability of the trees.

### Statistical analysis

A descriptive statistical analysis with absolute (n) and relative frequency of infection (%) was carried out, with a confidence level of 95%, using the free online tool WinEpi (Working in Epidemiology) and considering that the diagnostic techniques used were perfect (nPCR of the *SSUrDNA*). Odds ratio analysis was used to analyze the risk factors ‘sex’ and ‘*Leishmania* infection’ [25].

## Results

### *Detection and identification of* Leishmania *spp. in wildcats*

In this study, samples of the ear skin and spleen of 18 wildcats were analyzed, except for the spleen of two animals that were in poor condition when the cats were found (Table 1). None of the animals showed lesions comparable to those produced by *Leishmania* infections. Taking into account all animals positive for *Leishmania* PCR, 38.89% (16.37-61.41%, 95% confidence interval) were found to be infected. A nonsignificant higher proportion of infection was found in males (OR= 27.17, 95% CI 0.33, 2213.63), since only males were found infected, but we must consider that we obtained a smaller number of samples from females (n=3/18).

**Table 1.**
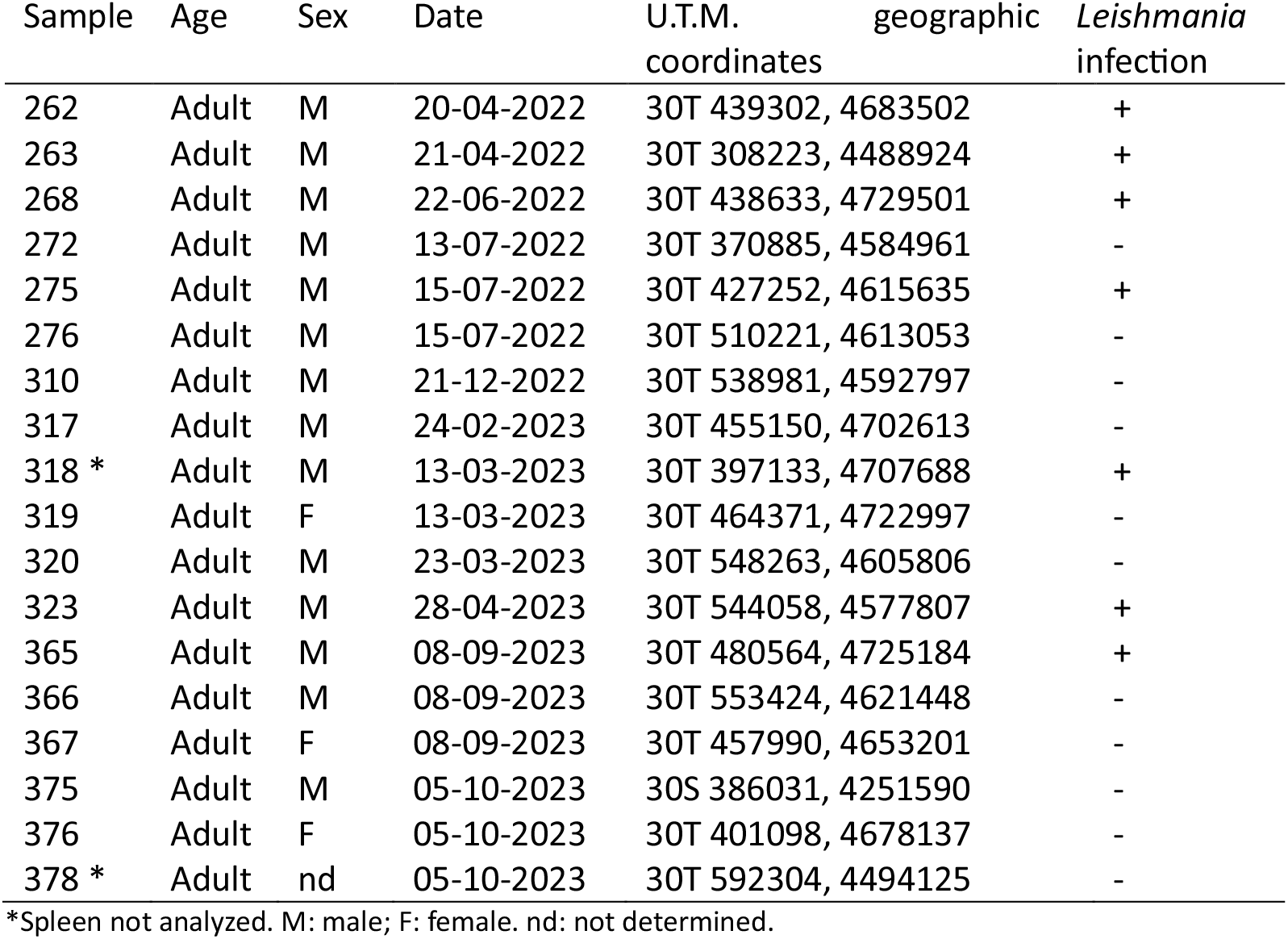
Age, sex, U.T.M. geographic coordinates, date of collection, and *Leishmania* infection of the studied wildcats.

All *Leishmania* spp. infected animals were found in areas where *P. sergenti*, the main vector of *L. tropica* was identified, except for one animal in which *L. infantum* was determined in both ear skin and spleen (Figure 1).

Seven of the 18 animals were found to be positive by *Leishmania* PCR for SSU-rDNA. All were positive on the skin of the ears and only two were also positive in the spleen samples (Table 2).

**Table 2.**
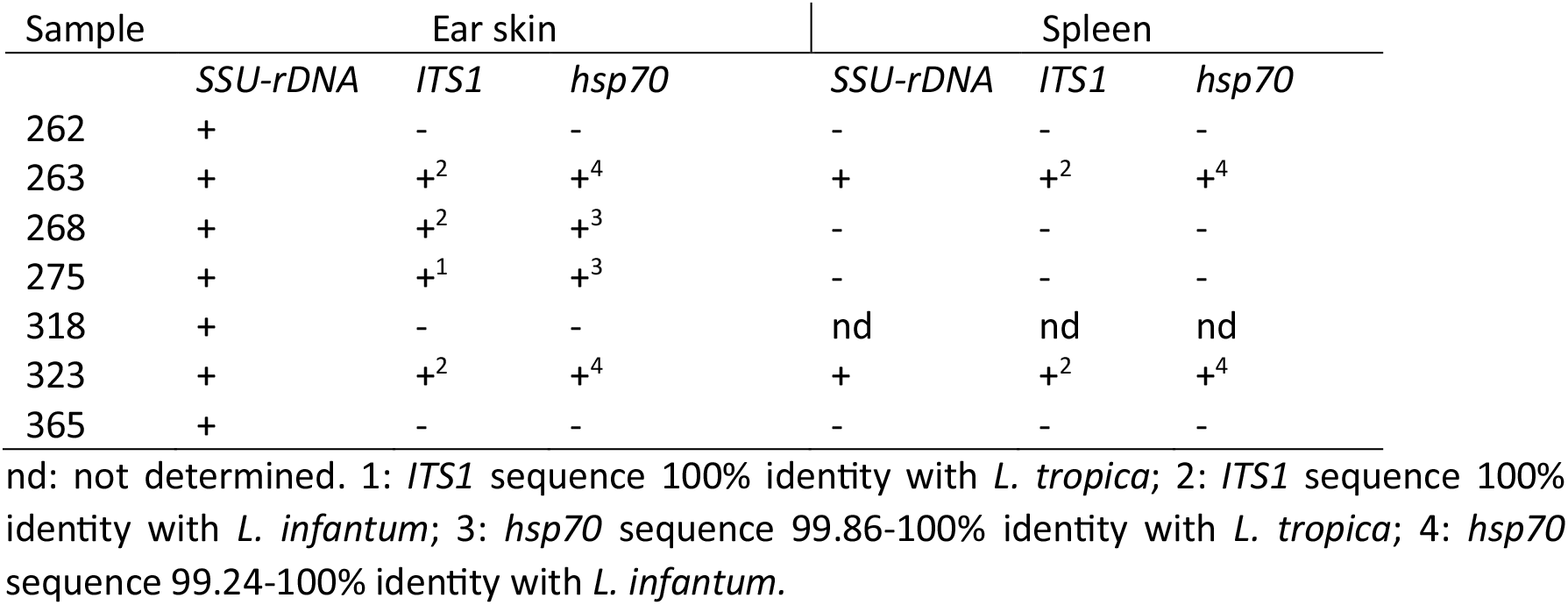
Results of endpoint PCR using samples from the ear skin and spleen and three targets for the amplification of *Leishmania* DNA. Sequence identities are identified with super index numbers (1-4).

### Sequences and GenBank accession numbers

The nucleotide sequences obtained in this study were deposited in the GenBank database under accession numbers PP389513-PP389521 for *SSU-rDNA*, PP177368-PP177373 for *ITS1*, and PP397157-PP397162 for *hsp70*. Sequences from *L. major* and *L. braziliensis* isolates were deposited in the GenBank database with accession numbers PP389522, PP389523, and PP389533.

The sequences obtained from *SSU-rDNA* had 295-327 bp and differ by only 1 or 2 nucleotide positions. Nine sequences were obtained from the animals analyzed, seven from the skin of the ears, and two from the spleen. All of them displayed 100% query coverage and 99.69-100% identity with other *Leishmania* spp. sequences available at GenBank, including *L. infantum* and *L. tropica*, among other species (acc. n MN757921, KF302745).

*ITS1* sequences of 236-247 bp were obtained from the ear skin of four animals (263, 268, 275, 323), two of which (263 and 323) also gave sequences when DNA from the spleen was used (Table 2). Three sequences displayed 100% query coverage and 100% identity with sequences from *L. infantum* (acc. n MT416168, samples from ear skin of animals 263, 268, and 323, and samples from spleen of animals 263 and 323), while one sequence (from wildcat 275 ear skin) displayed 100% query coverage and 100% identity with sequences from *L. tropica* (acc. n MH595857, isolate R24, Iran).

Sequences obtained from *hsp70* amplification yielded four sequences from the ear skin and two sequences from the spleen with 635-711 bp. Of these, four sequences showed 100% coverage and 99.24-100% identity with *L. infantum* (acc. n OR136937, strain MHOM/IT/99/ISS1898). Sequences 275 from the ear skin showed 100% coverage and 100% identity with *L. tropica* (acc. n MK335938, voucher ISS3183), while sequence obtained from the ear skin of 268 showed 100% coverage and 99.86 identity with *L. tropica* (acc. n LN907846, strain MHOM/IL/80/SINGER and others) and 99.72% identity with *L. donovani* (acc. n MH202961 and JX021427) (Table 2). Double peaks were observed at three and five positions of the sequences of the spleen of animal 323 and the skin of the ears of animal 263, respectively, indicating mixed sequences.

It is noteworthy to point out that *Leishmania* DNA was not detected by none of the three PCR methods in the spleens of cats with cutaneous infection with *L. tropica*. In one animal (275) the sequences from the *ITS1* and *hsp70* targets showed 100% identity with *L. tropica*. Two animals displayed sequences compatible with *L. infantum* on the skin of the ears and the spleen (263 and 323), and another wildcat was positive only on the skin of the ears with sequences compatible with *L. infantum* and *L. tropica* (268). Finally, in three animals (262, 318 and 365) the sequences could only be assigned to the *Leishmania* genus because they were only positive for *SSUrDNA* (Table 2).

### Phylogenetic trees

Phylogenetic analysis of the sequences shows strong support for the clusters found with the two targets, in agreement with the percentage of identity of the sequences obtained. *ITS1* sequences from this study were grouped with *L. infantum* (samples 263, 268 and 323 from the skin of the ears and samples 263 and 323 from the spleen) or *L. tropica* (sample 275) (Figure 2).

**Figure 2.**
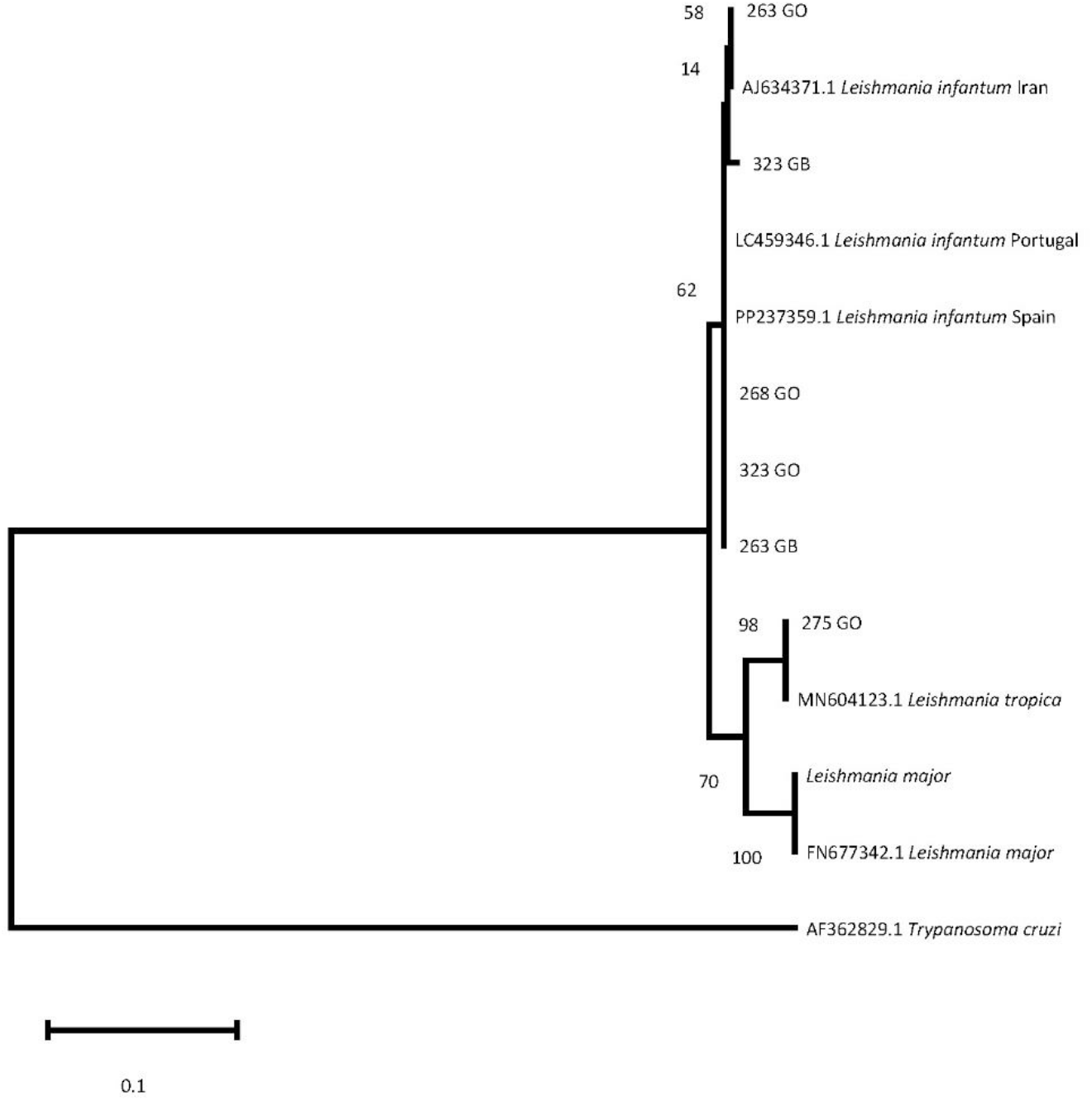
Phylogenetic tree of the *ITS1* region. The evolutionary history was inferred by using the maximum likelihood method and the Tamura-Nei model. The tree with the highest logarithmic likelihood (−841.79) is shown. The percentage of trees in which the associated taxa clustered together is shown next to the branches. The tree is drawn to scale, with branch lengths measured in the number of substitutions per site. This analysis involved 11 nucleotide sequences. There were a total of 292 positions in the final data set. GB: sample of the spleen. GO: sample from the skin of the ear.

In the phylogenetic tree generated with the *hsp70* sequences obtained in the present study, two of them (samples 268 and 275 from the ear skin) clustered with *L. tropica* and three of them (samples 263 and 323 from the ear skin and 263 spleen) clustered with *L. infantum* sequences recovered from GenBank (Figure 3). One of the *L. tropica* sequences was obtained from the same animal (275) that rendered the *L. tropica ITS1* sequence, while the other *L. tropica hsp70* sequence was obtained from an animal (268) that rendered the *L. infantum ITS1* sequence.

**Figure 3.**
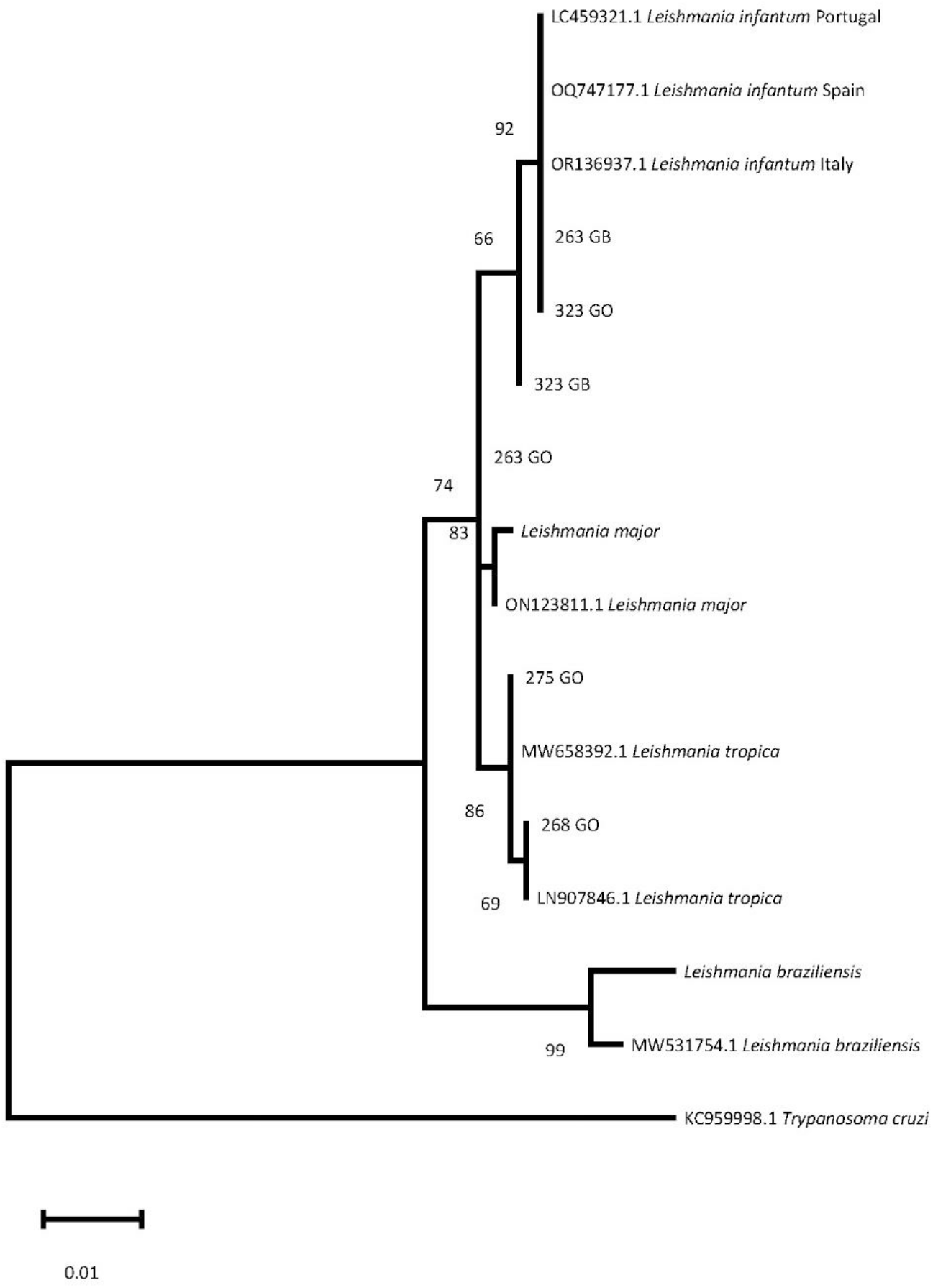
Phylogenetic tree of the *hsp70* region. The evolutionary history was inferred by using the maximum likelihood method and the Tamura-Nei model. The tree with the highest log-likelihood (−1328.37) is shown. The percentage of trees in which the associated taxa clustered together is shown next to the branches. The tree is drawn to scale, with branch lengths measured in the number of substitutions per site. This analysis involved 14 nucleotide sequences. There were a total of 635 positions in the final dataset. Evolutionary analyzes were carried out in MEGA11. **\***: Sequences obtained from positive samples from this study. GB: sample of the spleen. GO: sample from the skin of the ear.

## Discussion

In the present study, seven out of 18 wildcats (38.89%) were found to be positive for *Leishmania* infection, a value consistent with those found in previous studies [8,10] although lower than the infection prevalence reported by Oleaga et al. [9] with all three wildcats tested being positive for *Leishmania. L. infantum* is endemic in the Iberian Peninsula [12,13], but no infections of *L. tropica* in autochthonous species have been reported. Even considering the limitations of the study, including the low parasitic loads commonly found in wildlife and the number of animals, the information is of value, taking into account that the population of wildcats in Spain is considered relict and they are animals difficult to see unless they are sick or death, which is the case [26].

The risk of introduction of *L. tropica* in the Iberian Peninsula has been highlighted several times, considering the presence and distribution of the main vector, *P. sergenti*, and the identification in Spain of one genetic line of *P. sergenti* commonly found in Morocco [27,28] where CL by *L. tropica* is endemic [21,29,30]. Large areas in Europe are considered suitable for *L. tropica* [11] and hot spots for foci of *L. tropica* have been suggested [27]. In addition, the possibility of *L. tropica* transmission by *P. perniciosus* has been pointed out [31,32].

The most common targets used for the detection of *Leishmania* spp. infection are *SSU-rDNA* and kDNA, with similar values of sensitivity [6]. However, other targets are useful for phylogenetic studies and determination of *Leishmania* spp., such as *ITS1, cytB, g6pdh*, and *hsp70* [14,20– 22,29,33,34]. In the present study, *ITS1* and *hsp70* have been used to confirm the finding of *L. tropica* in autochthonous fauna, and sequences of *L. tropica* with *ITS1* and *hsp70* targets are reported. *L. tropica sequences* were detected in at least two wildcats positive for PCR with targets *hsp70* and / or ITS1, widely used for phylo-genetic studies. Our finding extends the reported presence of the species in Greece [15] and extends its distribution to the Iberian Peninsula for the first time.

The isolation of alive *Leishmania* was not feasible considering the source of material, but it is also noteworthy that in our study the skin samples of the ears showed more positive results than the spleen samples (n=7/18 *vs*. n=2/16). This finding is consistent with the fact that *L. tropica* infections are primarily associated with clinical forms of skin in humans, where the persistence of the parasite is close to the site of natural infections transmitted by the sandfly, although the parasite can also be found less frequently associated with VL, mainly in immune suppressed people [36]. Imported CL cases from *L. tropica* have been sporadically reported in Spain after travelling to Morocco [37] but no previous autochthonous cases have been reported in Spain or Portugal. Apparently, *L. tropica* is expanding its distribution and the main risk factor for the appearance of the disease is the presence of the vector [38]. It should be noted that some of the wildcats came from latitudes over 42º N (eg. animal 365, geographic coordinates UTM 30T 480564, 4725184) and during the last decade, *P. sergenti* has expanded to the northern areas of the Iberian Peninsula, according to ECDC maps of distribution [18].

In endemic areas, such as Israel, domestic cats appear to be more susceptible to infection with *L. tropica* than dogs [39]. A study of stray cats in Turkey showed that all samples with *ITS1* amplification were infected with *L. tropica*, while only one was co-infected with *L. infantum* [40]. Similarly, in the present study, while one animal showed infection by *L. tropica* only, another wildcat showed *hsp70* and *ITS1* sequences consistent with coinfection by *L. infantum* and *L. tropica* or by a hybrid of *L. infantum* and *L. tropica*. Mixed infections with more than one *Leishmania* species have also been reported in other endemic areas, hosts and *Leishmania* species, such as in Brazil (*L. infantum* and *L. braziliensis*) in one dog [41] and 8/30 dogs [42], rodents (10% of *L. infantum* and *L. braziliensis*) [(43)], hedgehogs (*L. major* and *L. infantum*) in 8/12 animals and another co-infected with *L. tropica* [(44), and human clinical cases [45,46] of the New World. The possibility of hybridization between *Leishmania* species cannot be ruled out, as *L. tropica* shows a higher rate of natural hybrid formation than other species [47]. Indeed, at least one natural hybrid of *L. donovani* and *L. aethiopica* has been described in Ethiopia employing *ITS1, hsp70* and cysteine proteinase B (*cpb*) targets [48]. The authors suggested that hybridization could take place in a vector permissive to both species, as it has been described in *P. pernicious* for *L. infantum* and *L. tropica* [31,32]. One hybrid of *L. infantum* and *L. major* was also described in a natural infection in Portugal [49]. However, to further investigate the existence of hybrids, isolates from infected animals are necessary [50,51], and in our study, they could not be obtained.

## Conclusions

This is the first report of *L. tropica* infection in autochthonous wildlife in the Iberian Peninsula, extending the presence of the species in continental Europe. The main vector of *L. tropica* has been identified in Spain during the last decade, and its distribution has increased northward each year. There are no conclusive data on the zoonotic transmission of *L. tropica* and the actual spread of the infection to other species is not yet known and should be investigated. Our results support the sentinel value of wild species in the detection of previously unreported infections. In addition, the presence of *L. tropica* in Spain should be communicated to health professionals, including physicians, dermatologists, and veterinarians, considering that *L tropica* can produce both CL and VL in humans and pets. Awareness of the possibility of *L. tropica* infections and *L. infantum*/*L. tropica* coinfections is necessary to ensure the appropriate diagnosis and management of clinical cases.

## Preprint article

A pre-print version of the manuscript with doi https://doi.org/10.1101/2024.03.16.585353 has been published on bioRxiv: https://biorxiv.org/cgi/content/short/2024.03.16.585353v1

## Data availability

Sequences obtained in the present study have been deposited in the GenBank database under accession numbers: PP389513-PP389521 for *SSU-rDNA*, PP177368-PP177373 for *ITS1*, and PP397157-PP397162 for *hsp70*. The sequences of the isolates of *L. major* and *L. braziliensis* are PP389522, PP389523, and PP389533.

## Ethical statement

The use of samples for parasitological investigations was approved by regional authorisation of the Junta de Castilla y León, reference: “AB/is”, File “AUES/CYL/001/2021” and of Castilla-La Mancha, reference: DGPFEN/SEN/avp_21_103_bis.

## Authors’ contributions

MMB, IAC, & MTG-M: conceptualization; IAC, PPM & MTG: methodology and formal analysis; MMB, PM & JC: collection of data and biological samples; IAC, MTG-M: writing the original draft; PMM, JC, IAC, MMB, MTG-M & JMA: interpretation of data; MMB & MTG-M: supervision; JMA, MMB & MTG-M: resources. All authors critically reviewed the manuscript, intellectually contributed to the content, and approved the final version.

## Acknowledgements

We deeply thank the cooperation of Junta de Castilla y León, Consejería de Fomento y Medio Ambiente, Dirección General de Patrimonio Natural y Política, and Castilla-La Mancha, Consejería de Desarrollo Sostenible, Dirección General de Medio Natural y Biodiversidad for permits to take samples of dead wildlife of the area.

